# Beta-driven closed-loop deep brain stimulation can compromise human motor behavior in Parkinson’s Disease

**DOI:** 10.1101/696385

**Authors:** Iñaki Iturrate, Stephanie Martin, Ricardo Chavarriaga, Bastien Orset, Robert Leeb, Aleksander Sobolewski, Etienne Pralong, Mayte Castro-Jiménez, David Benninger, Jocelyne Bloch, José del R. Millán

**Affiliations:** Defitech Chair in Brain-Machine Interface, Center for Neuroprosthetics, Ecole Polytechnique Fédérale de Lausanne, Switzerland; Helen Wills Neuroscience Institute, University of California, Berkeley, CA, USA; Clinical Neuroscience, University Hospital of Vaud (CHUV), Lausanne, Switzerland

## Abstract

Closed-loop or adaptive deep brain stimulation (DBS) for Parkinson’s Disease (PD) has shown comparable clinical improvements to continuous stimulation, yet with less stimulation times and side effects. In this form of control, stimulation is driven by pathological beta oscillations recorded from the subthalamic nucleus, which have been shown to correlate with PD motor symptoms. An important consideration is that beta activity is itself modulated during volitional movements, yet it is unknown the impact that these volitional modulations may have on the efficacy of closed-loop systems. Here, three PD patients performed a functional reaching task during closed-loop stimulation while we measured their motor behavior. Our results show that closed-loop stimulation can alter motor performance at distinct movement intervals. Of particular relevance, closed-loop DBS compromised behavior during the returning period by increasing the amount of submovements executed, and in turn delayed movement termination. Following these findings, we hypothesize that the use of machine learning decoding different movement intervals to fully switch off the stimulator may be beneficial, and present here an exemplary approach decoding the initiation of the movement returning interval above chance level. These findings highlight the importance of evaluating these systems during functional tasks, and the need of extracting more robust biomarkers encoding ongoing symptoms or tasks execution intervals.

## Introduction

Benefits of deep brain stimulation (DBS) for Parkinson’s disease (PD) are numerous (Limousin *et al.*, 2005; Deuschl *et al.*, 2006; Benabid *et al.*, 2009; Schuepbach *et al.*, 2013). However, several studies have reported side-effects affecting mood, behavior, moral competence, speech and personality (Klostermann *et al.*, 2008; Cyron, 2016). A promising approach to minimize those side-effects is to reduce the stimulation time using a closed-loop system, in which the stimulation parameters are adapted in real-time based on biomarkers reflecting the clinical state of the patient. One of such biomarkers for closed-loop deep brain stimulation (CLDBS) is the level of beta frequency (∼13-30 Hz) activity in the subthalamic nucleus (STN) of PD patients (Little *et al.*, 2013; Yamamoto *et al.*, 2013; Arlotti *et al.*, 2018; Velisar *et al.*, 2019), which has been shown to correlate with the severity of motor symptoms (Neumann *et al.*, 2016; Martin *et al.*, 2018).

Closed-loop systems in PD patients have shown enhanced or equal clinical improvements, reduced stimulation time, and reduced side effects compared to continuous stimulation (Little *et al.*, 2016). Despite these promising results, beta-driven CLDBS have only demonstrated their efficacy in treating motor symptoms as measured by clinical scores. Nevertheless, an important consideration is that beta activity is itself modulated during volitional movements (Litvak *et al.*, 2012), which may impact CLDBS efficacy during functional tasks. Indeed, a single non-human primate case study suggested that CLDBS performance compromised certain behavioral aspects during a motor task (Johnson *et al.*, 2016).

Here, we hypothesized that beta-driven CLDBS can also impact functional motor tasks performed by humans. For this, after ascertaining the advantage of CLDBS with seven human PD patients measured by clinical scores, three of these patients performed a motor reaching task during off, continuous and closed-loop subthalamic stimulation. With these data, we investigated how modulations in beta activity (desynchronization/synchronization) along with movements influenced the switching (on/off) of the stimulator, and in turn impacted motor behavior during the task. Our results highlight the importance of evaluating CLDBS not only using clinical scores but also during motor tasks, and the need for more robust biomarkers so that CLDBS remains efficient during functional tasks and activities of daily living.

## Methods

### Patients and data acquisition

Local field potentials (LFPs) were recorded between 36 and 48 hours after STN DBS bilateral implantation in seven PD patients. All patients volunteered and gave their informed consent. Experimental protocol was approved by the local ethical committee (CER-VD, Commission Cantonale d’Ethique VD de la recherche sur l’être humain). Long-acting dopaminergic medication was withdrawn 24 hours prior and short-acting medication was withdrawn 12 hours prior to off-medication pre-operative testing and to the DBS lead implantation procedure. All patients were operated by the same surgeon, providing homogeneity among the recordings. The awake surgical procedure stereotactically implanted a 3389 Medtronic electrode with 4 channels in the motor part of the subthalamic nucleus. Direct surgical planning was performed with the Medtronic stealthstation, based on a CT scan obtained with the CRW stereotactic frame, and fused with two 3 Tesla MRI sequences (MPRAGE with Gadolinium and space T2), obtained prior to surgery. After a first phase of microrecording, macrostimulation was performed under neurological clinical assessment. Adjustment of the electrode placement through a second track was performed to optimize the clinical response to stimulation. The electrode location was checked during the surgery with the O-Arm (Medtronic). An externalized extension cable was connected to the distal part of the electrode and tunneled posterior to the skin incision to record the brain activity for a period of 3 days.

### Closed-loop deep brain stimulation

We stimulated at 130 Hz unilaterally in the hemisphere contralateral to the most affected upper limb from the two contacts selected by the neurosurgeon. Our closed-loop system used a similar approach as in (Little *et al.*, 2013). First, a bipolar signal was derived from the two remaining contacts. Then, this signal was filtered within a beta sub-band free of stimulation artifacts (21-26 Hz), rectified and smoothed with a window size of 500 ms. Finally, the stimulation was triggered on/off based on a threshold, estimated as the 50th-percentile of the smoothed beta power during six minutes of resting-state data.

We validated our closed-loop approach in seven PD patients during six minutes of resting state by comparing their clinical scores during off, continuous, and closed-loop stimulation recorded in a random order (Fig. 1A). Clinical scores were assessed at the end of each condition by independent neurologists using the Unified Parkinson’s Disease Rating Scale (UPDRS) part III Motor Examination for resting tremor (item 20), rigidity (item 22) and bradykinesia (item 23). Prior to the recordings, the stimulation device was turned off for at least ten minutes.

**Figure 1.**
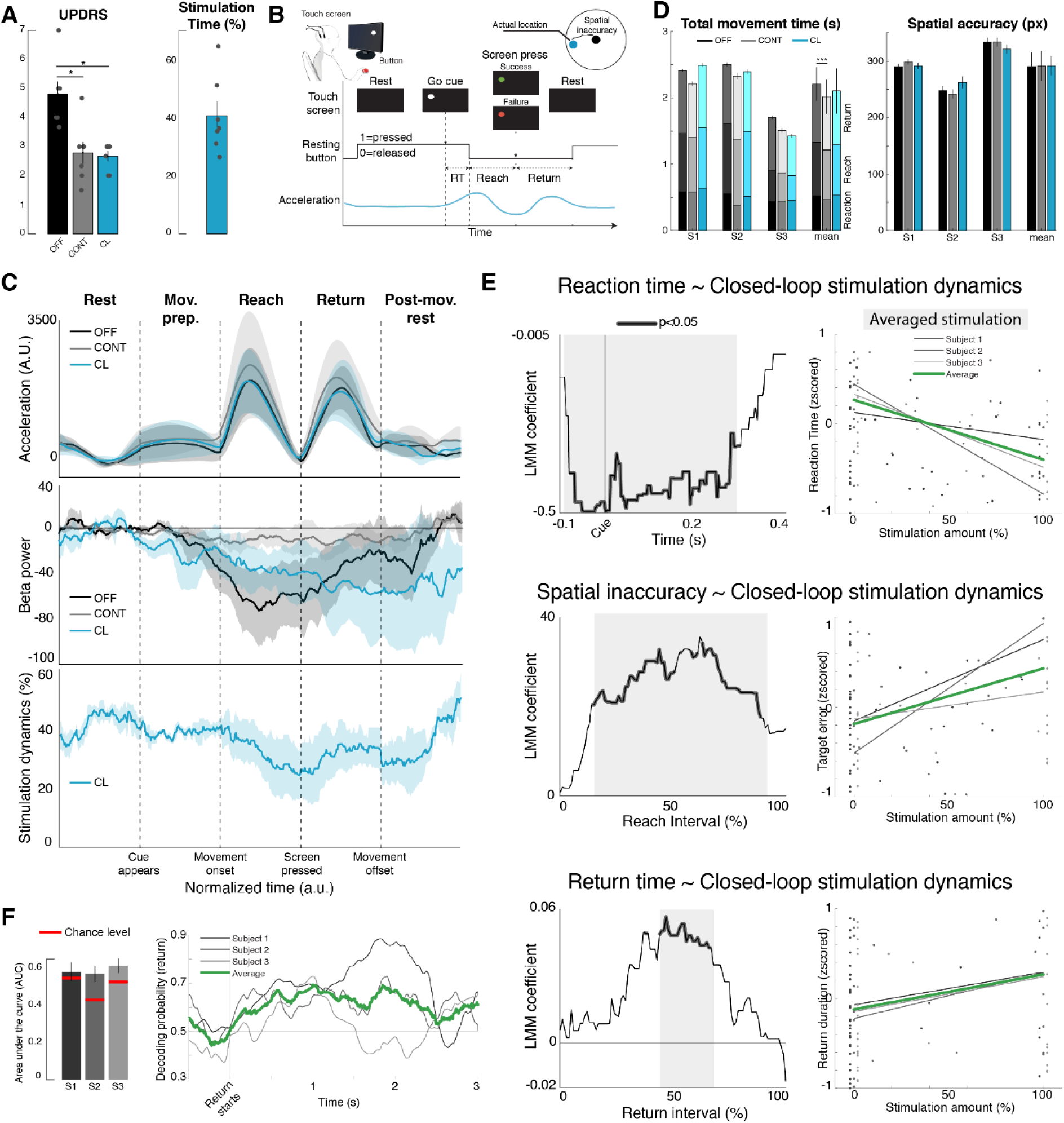
Impact of closed-loop DBS on movements. **(A)** Continuous and closed-loop DBS significantly improved motor symptoms (sum of UPDRS for bradykinesia, rigidity and tremor) when compared to not stimulating (N=7, p<0.05). Our closed-loop system stimulated only 40±13% of the time. **(B)** Patients performed a reaching task where they rested their most affected hand on a button while facing a tactile screen. When a cue appeared, patients had to reach and press the cue as fast and precisely as possible and come back to the resting position. We measured reaction, reach and return times; spatial inaccuracy and acceleration profiles. **(C)** Temporal interpolated evolution of acceleration modulus (top panel), ERD/ERS (middle) and stimulation dynamics during the closed-loop condition (bottom). **(D)** Movement times (left) and spatial inaccuracy (right) for the three conditions. The total movement time was significantly better in continuous with respect to off stimulation (N=264, p=10^−11^), but no differences were found between closed-loop and off conditions (p>0.4). **(E)** Time-resolved LMM of behavior as a function of closed-loop stimulation dynamics (reaction time, spatial inaccuracy and return time; for reach time see Fig S1). Left plots depict the LMM coefficients (N=133) and significant regions (shadowed regions). Right plots show the stimulation percentage averaged over the shadowed area (x-axis) versus behavior (y-axis). Stimulation around cue appearance led to faster reaction times, yet it significantly worsened spatial accuracy (i.e., spatial inaccuracy was larger) and returning times (p<0.05). **(F)** Successful decoding of the returning initiation is possible using LFP features. Left panel shows the area under the curve (AUC) together with their estimated decoding chance levels; right panel depicts the output probability of decoding the returning interval throughout time: probability increases right after the return period has started.

### Experimental paradigm

Three out of the seven patients performed a reaching motor task with their most affected hand in three stimulation conditions: off, continuous and closed-loop (Fig. 1B). A trial consisted of the following sequence: first, patients had to press a button placed in front of them, which marked the resting condition; then, a white cue of 400 pixel radius (approximately 3.8 cm) appeared at a random location on a touch-sensitive screen, and patients had to reach and touch the target with the index finger as fast and accurately as possible. In order to motivate participants, the cue turned green (red) if they pressed inside (outside) the cue area. Finally, patients returned to the resting position. Trials were repeated between 30 and 60 times per stimulation condition, depending on patient’s physical condition. During the task, we also recorded the hand kinematics using a 3-axis wireless accelerometer placed on the subject’s wrist (Shimmer sensing, sampling rate 50 Hz).

## Data analysis

We investigated how beta activity (21-26 Hz) and stimulation patterns (on, off, closed-loop) related to movement kinematics and behavior. We first extracted the hand kinematics by filtering each accelerometer dimension with a non-causal, fourth-order Butterworth filter within 0.5 and 6 Hz and calculated the modulus.

Then, we epoched beta activity, stimulation state and movement kinematics for each trial. In order to investigate the different phases of movement, we split the trials into five distinct *movement intervals*: pre-movement rest, movement preparation, reach, return, and post-movement rest (Fig. 1B,C). We resampled each interval to the percentage of interval completed (0% represents the beginning of the interval, 100% represents the end of the interval) in order to compare movements that had different time lengths across trials and patients.

We evaluated the impact of each stimulation condition (off, continuous, closed-loop) on five behavioral metrics: reaction time, reach time, return time, total movement time and spatial inaccuracy (measured as the distance in pixels between the target and the reached position). For each behavioral metric, we compared off vs continuous and off vs closed-loop using linear mixed models (LMMs). LMMs are an extension of linear models used when data are not independent (e.g., a patient is altogether faster than another patient) and thus allow using all individual trials from all patients (N=264). We defined the LMM as follows:

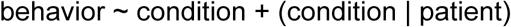

The aim of this LMM was to find whether continuous or closed-loop stimulation had any significant impact on behavior relative to off stimulation (term *condition*). By controlling for random slopes and intercepts for each patient (term *(condition | patient)*), we ensured that any effect detected as significant was shared across all patients. The obtained p-values in these analyses were corrected for multiple comparisons using the false discovery rate (FDR) (Storey, 2002).

In the closed-loop condition, the stimulation state (on/off) varied over time. We named these variations of stimulation *closed-loop stimulation dynamics*. We hypothesized that variations at specific time points during movements may have differential outcomes on behavior. Thus, we investigated the impact of stimulation during closed-loop throughout the executed movement (N=133) with another LMM:

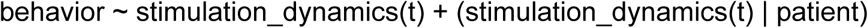

For this analysis, we were interested in the coefficients (also termed slopes) estimated for the stimulation dynamics at each time point *t* throughout the movements. These coefficients define how the dynamics relate to behavior: a positive (negative) coefficient indicates a direct (inverse) relationship between stimulation dynamics and behavior, and larger coefficients indicate stronger relation between the amount of stimulation and the behavioral metric. Such coefficients are accompanied by their associated p-values, which indicate whether the effect is significant or not. We report both the coefficients and their associated p-values FDR corrected for multiple comparisons.

If significant differences were found in reaching or returning times behavioral metrics, follow-up LMM statistics on acceleration metrics were performed to discern the reason behind differences in movement times. First, we studied whether stimulation during closed-loop caused any change in peak speed during the reach/return movement (obtained as the maximum value of acceleration), following previous works (Johnson *et al.*, 2016). Second, we studied whether stimulation altered the total number of submovements executed during a reach/return, a metric that has been shown to be impacted in Parkinsonian patients (Dounskaia *et al.*, 2009). The number of submovements of each trial was obtained with a submovement decomposition algorithm using gradient descent (Gowda *et al.*, 2015). Finally, the associated p-values were FDR corrected.

### Decoding approach

Our results show that stimulation compromised the behavior during the returning period (see Results section). One plausible solution to solve this is to build more intelligent CLDBS approaches able to detect different movement intervals and acting accordingly (e.g., switching off the stimulation during the returning period). Here, we exemplified this approach by evaluating the ability to decode reach and return intervals from the recorded LFP in the STN.

For this, we employed a similar approach to (Orset *et al.*, 2019). We first extracted as features the beta power from both hemispheres within a 0.5 second window, leading to an initial set of 512 features. To train the decoder, an automatic algorithm chose the 10 best features with inner cross-validation based on their discriminability scores. Feature vectors in the training dataset were z-scored and used to build a diagonal linear discriminant (DLDA). To test the decoder, we extracted the features using a sliding window with shifts of 12.5 ms, in time intervals surrounding the returning period, [-0.5, 0] s and [0, 0.5] s, with respect to the time when the patients touched the screen (t=0). The decoding performance was evaluated using 5-fold cross-validation, from which we computed the area under the curve (AUC). For each patient, we also estimated the chance level by permuting the labels 500 times and followed the same train/test procedure.

### Data availability

The data that support the findings of this study are available from the corresponding author upon reasonable request.

## Results

We evaluated efficacy of our closed-loop system in seven PD patients measured from their UPDRS clinical scores from bradykinesia, rigidity and tremor. Clinical symptoms significantly improved during continuous (p=0.02; signed rank test; FDR-corrected) and closed-loop stimulation (p=0.02; signed rank test; FDR-corrected) compared to off stimulation (Fig. 1A; left panel). Both, continuous and closed-loop stimulation performed equally well (no differences observed; p=1; signed rank test), yet CLDBS stimulated only 40±13% of the time (Fig. 1A, right). We found no significant differences in the total stimulation time during rest state versus the stimulation time during UPDRS evaluation windows (43±15% and 41±14% resp., p=0.47, signed-rank test). These clinical outcomes are in agreement with previous reports on symptoms improvements using both continuous and closed-loop DBS (Little *et al.*, 2013).

Three of these patients performed a reaching task with their most affected hand during off, continuous and closed-loop stimulation (Fig. 1B). Acceleration profiles showed typical movement dynamics, with peak accelerations for all conditions at 40% and 50% during reaching and returning, respectively (Fig. 1C, top). We observed a beta desynchronization and resynchronization for the three conditions, although during continuous the relative modulations were smaller (Fig. 1C, middle). Due to these beta modulations, stimulation during the closed-loop condition was not stationary but dynamically fluctuated throughout the movement intervals, decreasing during movement onset and increasing again during the returning period and rest intervals (for each patient, 5-67%, 30-53%, 24-51% of minimum and maximum stimulation times within movements, computed as the percentage of trials stimulated, Fig. 1C, bottom). In sum, beta modulations during volitional movements interfered with the closed-loop control, which in turn may affect successful desynchronization/resynchronization and its associated behavior.

When comparing overall behavioral outcomes of closed-loop and continuous stimulation against off, behavior during closed-loop stimulation was not significantly different than off stimulation for any metric (N=264, p>0.4). Conversely, continuous stimulation led to faster reaching times (p=0.0005, LMM, FDR-corrected) and total movement times (p=10^−11^) compared to off stimulation (Fig. 1D), with the rest of behavioral metrics not different between conditions (p>0.4). This first comparison indeed suggested that closed-loop stimulation may not be as effective as continuous stimulation in improving behavior during a motor task.

We hypothesized that, for the closed-loop stimulation condition, stimulating at specific time points during movements may have differential outcomes on behavior. To prove this hypothesis, we developed a time-resolved analysis which tried to predict behavior from the closed-loop stimulation dynamics, by using one LMM per time point during the movement. With this analysis, we found that stimulating around the cue appearance period led to significantly faster reaction times (N=133, [-0.1, 0.3] s, FDR-corrected p<0.05, Fig. 1E, top). This effect was present for the three patients (see average fittings over the significant period, Fig 1E, top, right). While CLDBS during the reaching interval did not influence reaching times (p>0.22, Fig. S1), spatial accuracy significantly worsened with stimulation for all three patients (i.e., spatial inaccuracy was larger, [14, 89]% of the executed movement, p<0.05, Fig. 1E, center). Finally, stimulation also slowed movements during return intervals for the three patients (p<0.05, [44, 67]% of the movement, Fig. 1E, bottom), yet this slowing was not due to speed differences (p>0.1). Contrarily, the slowing was due to a greater number of submovements when stimulation was applied at around half of the returning period (p<0.05, [36, 60]% of the movement, Fig. S1). In sum, stimulation led to a greater amount of submovements, which in turn led to longer returning times.

These results show that closed-loop stimulation can compromise behavior. One plausible solution to solve this limitation is to build smarter CLDBS approaches able to detect different movement types and switching the stimulator on/off accordingly. Here, we exemplified this approach by evaluating the ability to decode the moment when the reaching interval finished, and the returning interval started using the recorded LFP-STN signals. Reach vs return intervals were successfully decoded during closed-loop stimulation (decoders used bilateral beta features, area under the curve, five-fold cross-validation AUC= 58, 57 and 61% for patient 1, 2, 3, respectively; Fig. 1F, left). This returning phase was decoded significantly above chance as soon as 10 ms after it started (sliding window approach, decoding probability of return > 0.5, Fig. 1F, right).

## Discussion

Closed-loop DBS efficacy as measured by clinical scores is comparable to that of continuous stimulation (Little *et al.*, 2013), yet with less stimulation times and side effects (Little *et al.*, 2016). Although promising, a recent single case, non-human primate study has questioned its efficacy during motor tasks (Johnson *et al.*, 2016). Here, we show how closed-loop DBS can impact behavioral performances during a reaching task performed by three human PD patients.

While continuous stimulation significantly improved the total movement time with respect to no stimulation, closed-loop DBS did not present such improvement. Contrary to previous works, continuous DBS did not lead to any significant differences in reaction times nor spatial accuracies. The reason for this absence is a constraint of our statistical model, which was designed to detect as significant only those effects that were consistent across the three subjects. Indeed, we found significant differences on single-subject analysis for reaction times and spatial accuracies in two out of three patients.

By using a novel, time-resolved analysis during closed-loop stimulation, we were able to pinpoint with a high time resolution the impact of stimulation in behavior throughout the movement, and only if the effect was consistent across the three patients to counteract the low sampling size. Of note, thanks to the use of linear mixed models, we can exploit single trials to have a sufficient number of data points to run the analysis. In this regard, stimulation surrounding the cue onset led to faster reaction times, in line with recent results proposing that beta oscillations delay movement onset (Khanna and Carmena, 2017); while it had a negative effect on spatial accuracy, an effect that has been elsewhere discussed (Agostino *et al.*, 2008).

The novel finding of this work lies on the fact that CLDBS also compromised behavior during the returning period. In this work, stimulation during the returning interval correlated with the execution of additional corrective submovements, potentially because it prevented beta resynchronization. As a consequence, patients needed more time to finalize the movement, in line with results suggesting that PD causes deficiency in movement termination due to gross submovements (Dounskaia *et al.*, 2009). Of note, our patients were not tremor dominant, which rules out the possibility that these submovements were tremor-driven.

A plausible explanation for this result is the nature of beta modulations during this returning period. While it is known that beta resynchronizes *after* a movement, there seems to be no consensus on the existence of beta resynchronization *during* certain movements intervals. As a supporting finding to ours, previous works have shown that goal-oriented movements lead to beta desynchronizations, but beta modulations during self-paced movements are almost non-existent (Bichsel *et al.*, 2018). In our case, beta resynchronizes during the returning period, coinciding to the switching from a cued movement (reaching) to a non-cued one (returning), as shown in the beta modulations during the off condition. Indeed, we have also found such an effect in cortical M1 modulations of a healthy population, see Fig S2 and (Iturrate *et al.*, 2018). We believe that, similarly to the fact that beta desynchronization is needed for movement initiation, our results suggest the functional relevance of beta resynchronization as a driver of movement termination, and that closed-loop DBS compromised such termination.

Our results point out the inability of current CLDBS approaches to differentiate between volitional desynchronization/resynchronization beta modulations as those present during movements; and pathological beta power increases. Although preliminary due to the low population, our findings are consistent across the three subjects; and emphasize the need of further evaluation of CLDBS approaches in functional tasks, and to identify more robust biomarkers to trigger stimulation.

We hypothesized that machine learning may also be a promising path to alleviate these issues, mainly by detecting different movement intervals or conditions where stimulation may compromise performance. Here, we focused on trying to improve the returning behavior, and exemplified it with a decoder exploiting beta features able to significantly detect when the returning interval started, which could in turn be used to switch off the stimulation and override typical beta biomarkers. Alternatively, by exploiting a larger set of features coming from different LFP-STN oscillations, one could aim at dissociating pathological from physiological modulations; or detecting different stages during typical functional movements.

Added to the already promising results of current CLDBS approaches, we advocate that the success of new studies will be largely determined by their performance during functional tasks. Such studies will greatly impact new generations of CLDBS for PD patients, robust enough to work in out-of-the-clinic scenarios, and more importantly, without compromising their activities of daily living.

## Abbreviations

DBS: deep brain stimulation
LFP: local field potentials
PD: Parkinson’s disease
STN: subthalamic nucleus
LMM: linear mixed model
UPDRS: Unified Parkinson’s Disease rating scale
CLDBS: closed-loop deep brain stimulation
AUC: area under the receiver-operating characteristic curve

## Acknowledgement

We thank Michael Pereira, Elvira Pirondini, Mark Hallett, Leonardo G Cohen and Leonardo Claudino for their comments on the manuscript and help on the recordings. B.O. and J.d.R.M. were supported by the Swiss National Centres of Competence in Research (NCCR) Robotics. I.I. also acknowledges support from the ‘EPFL Fellows’ fellowship program co-funded by Marie Curie, FP7 Grant agreement no. 291771. DB is supported by the Foundation Parkinson Switzerland, the Swiss Dystonia Society and the Swiss Accident Insurance Fund, and holds the Baasch-Medicus Prize.

## Author contributions

I.I. Research Project: Conception, Organization and Execution. Statistical Analysis: Design and Execution. Manuscript: Writing of the first draft. S.M. Research Project: Conception & Execution. Statistical Analysis: Design & Execution. Manuscript: Writing of the first draft. R.C. Research Project: Conception, Organization and Execution. Manuscript: Review and Critique. B.O. Statistical Analysis. R.L. Research Project: Conception, Organization and Execution. Manuscript: Review and Critique. A.S. Research Project: Conception, Organization and Execution. Manuscript: Review and Critique. E.P. Research Project: Organization. Manuscript: Review and Critique. M.C.-J. Research Project: Organization. Manuscript: Review and Critique. D.B. Manuscript: Review and Critique. J.B. Research Project: Conception, Organization and Execution. Manuscript: Review and Critique. JdR.M. Research Project: Conception and Organization. Manuscript: Review and Critique.

