## Supplementary Information for "Beta-driven closed-loop deep brain stimulation can compromise human motor behavior in Parkinson’s Disease"

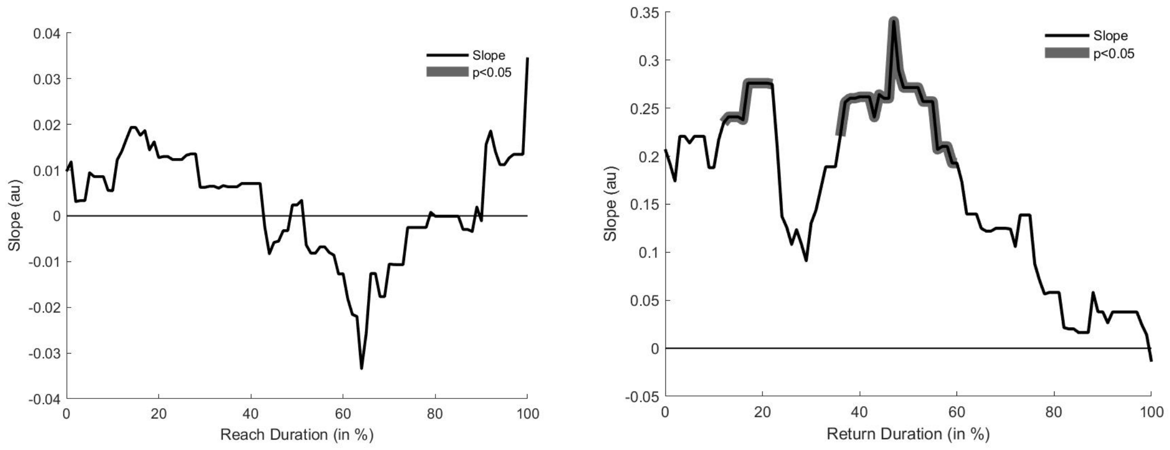


**Fig S1.** LMM time-resolved analysis for reaching time (left) and number of submovements during the returning period (right).

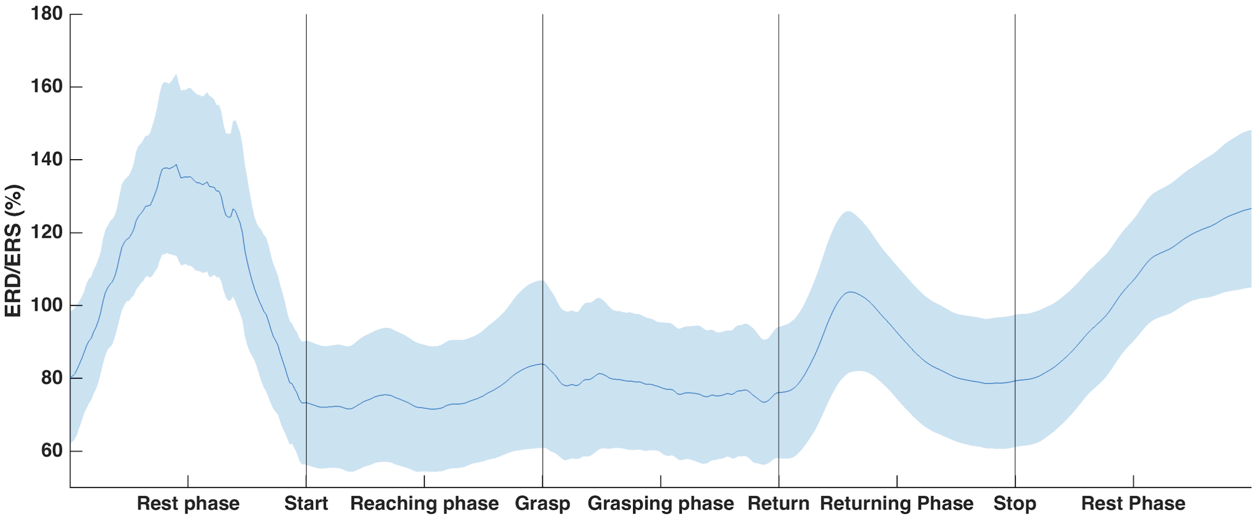


**Fig S2. Beta modulations during a reaching and grasping task.** In this experiments, nine healthy volunteers performed a self-paced reaching and grasping task while their electroencephalographic activity was recorded from 64 channels. During the reaching and grasping phases, a consistent beta desynchronization was observed over the contralateral motor cortex. During the returning phase, a beta resynchronization could be observed, with a larger, slower resynchronization during the post-movement resting phase, similar to the findings reported in this work. Results in this figure are interpolated, similar to those reported in Figure 1C of the main manuscript. For more information, see Iturrate *et al.* (2018) (reference in the main manuscript).
